# Incorporating fire-smartness into agricultural policies minimises suppression costs and ecosystem services damages from wildfires

**DOI:** 10.1101/2023.01.20.524753

**Authors:** Judit Lecina-Diaz, María-Luisa Chas-Amil, Núria Aquilué, Ângelo Sil, Lluís Brotons, Adrián Regos, Julia Touza

## Abstract

Global climate warming is expected to increase wildfire hazard in many regions of the world. In southern Europe, land abandonment and an unbalanced investment toward fire suppression instead of prevention has gradually increased wildfire risk, which calls for a paradigm change in fire management policies. Here we combined scenario analysis, fire landscape modelling, and economic tools to identify which land-use policies would minimise the expected wildfire-related losses in a representative mountainous area of the northwestern Iberian Peninsula (the Transboundary Biosphere Reserve ‘Gerês-Xurés’, between Spain and Portugal). To do so, we applied the least-cost-plus-net-value-change approach and estimated net changes in wildfire damages based on their implications for the ecosystem services that affect financial returns to landowners in the study area (i.e. agriculture, pasture, and timber) and the wider economic benefits (i.e. recreation and climate regulation) for the 2010-2050 period. Four land-use scenarios were considered: (1) Business as Usual (BAU); (2) fire-smart, fostering more fire-resistant (less flammable) and/or fire-resilient landscapes (fire-smart); (3) High Nature Value farmlands (HNVf), wherein the abandonment of extensive agriculture is reversed; and (4) a combination of HNVf and fire-smart. We found the highest net value change (i.e. the difference between damages and avoided damages) in BAU for timber and pasture provision, and in fire-smart for recreation and climate regulation. HNVf was the best for suppression cost savings, but it generated the lowest expected present value for climate regulation. In fact, the best scenarios related to fire suppression are HNVf and HNVf combined with fire-smart, which also generate the lowest net value change plus net suppression costs in the entire study area (i.e. considering all ecosystem services damages and suppression costs). Therefore, reverting land abandonment through recultivation and promoting fire-resistant tree species is the most efficient way to reduce wildfire hazard. In this sense, payments for ecosystem services should reward farmers for their role in wildfire prevention. This study improves the understanding of the financial and societal benefits derived from reducing fire suppression spending and ecosystem services damage by undertaking fire-smart land-use strategies, which can be essential to enhance local stakeholders’ support for wildfire prevention policies.

**Highlights:**

- Land-use changes impact wildfire ecosystem services (ES) damages and suppression costs
- Promoting agriculture generates significant suppression cost savings
- Agriculture + fire-resistant forests is the best to reduce wildfire ES damages
- Land-use policies should balance trade-offs between climate and wildfire regulation
- Payments for ES should reward farmers for their role in wildfire prevention

## Introduction

Global climate warming is expected to increase wildfire hazard in many regions worldwide, leading to large-scale changes in fire regimes (Abatzoglou and Williams, 2016; IPCC, 2018). In southern Europe, land abandonment has been a dominant factor contributing to wildfire risk by increasing fuel accumulation and continuity (Moreira et al., 2011). Fire management is primarily centred on suppression orientation measures (Fernandes, 2013). Billions of euros have been invested in fire suppression, with approximately 2,500 million € annually in the Mediterranean basin in recent years (Verkerk et al., 2018). This focus on fire suppression instead of strategic wildfire management (Castellnou et al., 2019) has contributed further to the large-scale accumulation of fuel that combined with rising temperatures and more frequent and severe droughts, results often in more severe and larger fires with high economic losses (San-Miguel-Ayanz et al., 2013, see ‘fire-fighting trap’ in Moreira et al., 2020). For example, large fires in 2005 and 2017 in Portugal caused losses of 800 million € and 1.5 million €, respectively (San-Miguel-Ayanz et al., 2020, 2013); and wildfires in black pine afforestation in northern Spain were valued under 124 €/ha (Alcasena et al., 2016).

There is, therefore, a demand for a paradigm change in fire management policies from suppression to prevention, addressing the interlinked social, ecological, and economic systems behind the ignition and spread of wildfires (Moritz et al., 2014; Moreira et al., 2020), especially in complex socio-ecological systems such as the abandoned landscapes of southern Europe. Fire-smart management has been proposed as “an integrated approach primarily based on fuel treatments through which the socio-economic impacts of fire are minimised while its ecological benefits are maximised” (see Hirsch et al. 2001; Fernandes, 2013). Fire-smart management (e.g. conversion from fast-growing tree plantations to more fire-resistant woodlands) was predicted to have positive effects on climate regulation while also contributing to fire regulation, i.e. the capability of landscapes to regulate spatiotemporal properties and characteristics of fire through the control of key factors that determine how fire behaves and its effects (Pettorelli et al., 2018; Sil et al., 2019). In that sense, agricultural policies have been claimed as a ‘fires-smart’ solution since they can contribute to the increase of landscape heterogeneity, reducing fire spread and lowering the continuity of fuel, accumulated due to rural abandonment and fire exclusion policies (Moreira and Pe’er, 2018; Pais et al., 2020). However, in Europe, the Common Agricultural Policy (CAP) failed to reverse rural abandonment and preserve biodiversity (Pe’er et al., 2020). The new CAP offers an opportunity to secure the future of agriculture and forestry sectors as well as to achieve the objectives of the European Green Deal, especially in mountainous rural landscapes where low-intensity agricultural and livestock activities are often associated with areas rich in biodiversity (also known as High Nature Value farmlands, HNVf) (Lomba et al., 2015). There is therefore an increasing need to incorporate fire-smart management into the upcoming European land-use policies to reduce wildfire hazard while preserving biodiversity and regulating climate (Fernandes, 2022; Regos, 2022).

Agricultural and forest landscapes provide a suite of ecosystem services; thus, any preventing wildfire management strategy based on land-use changes would inevitably lead to ecosystem service trade-offs (Mercer et al., 2008). Therefore, a comparison of the long-run net expected benefits of fire-smart strategies to reduce wildfire hazard is essential not only to support decision-making but also to assess the economic efficiency of government budget spending needed to support such land-use policies (i.e. ensuring that those expenditures pay off and are therefore justifiable). However, the application of economic analysis to assess the effectiveness of wildfire prevention strategies has been limited to fuel management through prescribed burning or mechanical fuel treatment (Hesseln, 2000; Mercer et al., 2007; Prestemon et al., 2012). In this regard, some studies have taken a public policy evaluation perspective and conducted an ex-post assessment of effectiveness measures using biophysical indicators such as the occurrence and intensity of wildfires (Butry et al., 2010; Lydersen et al., 2017). Ex-ante analysis, such as cost-benefit or cost-effectiveness analysis, is advisable when a manager is interested in choosing among alternative wildfire management actions that provide the highest societal net benefits (Butry and Prestemon, 2019). Cost-effectiveness analyses are used more frequently in the literature because the expected benefits of avoiding fires do not need to be expressed in monetary terms. For example, Penman et al. (2014) and Elia et al. (2016) evaluated the cost-effectiveness of fuel load removal, which reduced the likelihood of wildfire damage to houses. Similarly, economic studies have focused on determining the optimal investment level for wildfire management. Donovan and Rideout (2003) provided a rigorous analysis of the “cost-plus-net-value-change” (C+NVC) criteria to identify the most efficient level of fire management expenditure, where the decision-maker is set to minimise all management-related expenditures and wildfire damages. The recent work of Florec et al. (2020) used these criteria to evaluate the efficiency of alternative arrangements of prescribed burning treatments in the landscape. In an earlier study, Prestemon et al. (2012) questioned how much forestland can be mechanically treated with positive long-run net benefits, including benefits derived from timber revenues from treatment, and those derived from a reduction in wildfire occurrence (e.g. avoided timber losses, reduced suppression costs, and reduced property damage). Their results illustrated that, unless timber sales from mechanical treatment are allowed, such a prevention strategy will not generate net societal benefits. Moreover, ecosystem assessments that economically evaluate the trade-offs between the different impacts of land-use change policy interventions have recently been used in wildfire literature, such as the study by Raviv et al. (2021), which estimates the change in ecosystem services values affected by wildfires and consequent land-cover changes. However, there is little understanding of how policies that promote land-use changes (e.g. agricultural conversion or forest-type conversion), i.e. fire-smart management strategies, can contribute to reducing wildfire economic losses.

To this end, we combined scenario analysis, fire landscape modelling, and economic tools to evaluate the economic efficiency of alternative land-use policies as a wildfire prevention strategy. The question addressed here is which land-use policies would minimise the expected wildfire-related losses. Four competing land-use scenarios were evaluated: (1) a Business as Usual (BAU) scenario, which represents the long-standing rural abandonment; (2) a scenario that assumes a gradual conversion from forestry plantations to native oak woodlands (fire-smart); (3) a contrasting scenario that envisages a policy that aims to promote extensive agriculture and livestock activities (HNVf); and (4) an integrative scenario where policies ensuring HNV farmlands are combined with large fire-smart forest conversions. Our fire-landscape modelling approach allowed us to explore the spatial interactions between fire ignition, spread and suppression, and land-use changes at the landscape level over time. This allows us to evaluate the expected gains of land-use interventions in reducing burned areas due to the suppression capabilities of the simulated landscape policies. We then estimate the “cost-plus-net-value-change” in the form of the present value of future net reductions in suppression costs and wildfire damages to ecosystem services. Impacted ecosystem services flows included those that affect landowners’ private returns, e.g. from the provision of crops, and to the wider society such as recreation and climate regulation. Therefore, this study improves the understanding of the financial and societal benefits derived from reducing fire suppression spending and ecosystem services damage by undertaking fire-smart land-use strategies, which can be essential to enhance local stakeholders’ support for wildfire prevention policies. The case study is the Transboundary Biosphere Reserve Gerês-Xurés (Spain-Portugal), where the current abandonment of traditional and livestock activities in this area is representative of the socio-demographic and wildfire management challenges in the mountainous areas of the northwestern Iberian Peninsula.

## 2. Material and methods

We followed the cost-plus-net-value change approach, which is widely used for economic evaluation of fire management programs as a conceptual framework to evaluate alternative land-use policies (e.g. Florec et al., 2020; Houtman et al., 2013; Rodríguez y Silva and González-Cabán, 2010). This approach assumes that wildfire managers follow a cost-minimising behaviour, and the most economically efficient land-use change scenario would be the one that minimises the management cost plus the net-wildfire-related losses (value). Our evaluation criterion includes only suppression costs as a management cost. The cost of implementing land-use interventions was not considered here because most of the land-use changes simulated were through natural regeneration (e.g. natural succession, vegetation encroachment, or forest expansion due to abandonment), with some exceptions as HNVf strategies which promote cropland expansion. We modelled net-value-change (NVC) as net wildfire damages that reflect the effect on natural asset values. Therefore, we evaluated the net changes in ecosystem services flows associated with each land-use scenario for all years simulated. Thus, our analysis relies on the estimation of the effects of simulated land-use changes on wildfire ignition, spread, extinction, and the resulting burned areas per land cover. Therefore, using a process-based model developed for the study area (Pais et al, 2020), we simulated the spatial interaction of fire ignition, spread and suppression, and vegetation dynamics in the case study area under different land-use scenarios over a 40-year period (2010-2050). Our economic estimation of the present value of net wildfire damages from changes in ecosystem services flows uses the land cover map output from the REMAINS model (i.e. the burned and avoided area, identified separately for each land cover) as inputs. The counterfactual for all scenarios was the expected burned area, assuming that fire ignition and spread followed a historical trend (1987-2010) without suppression. This counterfactual allows us to compare the capacity of the simulated landscapes policies to suppress fire, and achieve benefits in terms of reduction in burned area. We detail below the case study area, scenarios simulated, fire landscape, and economic framework for the analysis.

### 2.1. Study area

The Gerês-Xurés transboundary Biosphere Reserve is located between 41° 35’ and 42° 10’ latitude North and 7° 35’ and 8° 31’ longitude west, covering an area of 2,760 km^2^ (71% in Portugal and 29% in Spain) (Figure 1). It was established in 2009 and includes three EU Natura 2000 sites and two nationally designated protected areas (the Peneda-Gerês National Park in Portugal and the Baixa Limia-Serra do Xurés Natural Park in Spain). Ranging in elevation from 15 to 1,545 m.a.s.l., it includes deep valleys, plains, and steep slopes (Regos et al., 2015). It is located at the transition between the Mediterranean and Eurosiberian biogeographic zones, with a climate mainly Atlantic (monthly average temperature below 22 °C; Kottek et al. 2006). The predominant landscape is dominated by heathlands, as well as fragmented forests of deciduous trees (mostly *Quercus robur* and *Q. pyrenaica*) and conifers (*Pinus sylvestris* and *P. pinaster*) (Figure 1). Rural abandonment has been a common trend in the area during the last century (Macedo et al., 2009; Regos et al., 2015). The population decreased by 17.8% from 2011 until 2021 (Martins, 2022), with a current population density of 29.4 inhabitants km^2^. Frequent human-caused wildfires are common in the study area (Calviño-Cancela et al., 2016; Chas-Amil et al., 2015, 2010). Consequently, there were a large number of fires and a total burned area (i.e. 12,755 fires between 1983 and 2010 burning a total of 195,000 ha, Regos et al., 2015).

**Figure 1.**
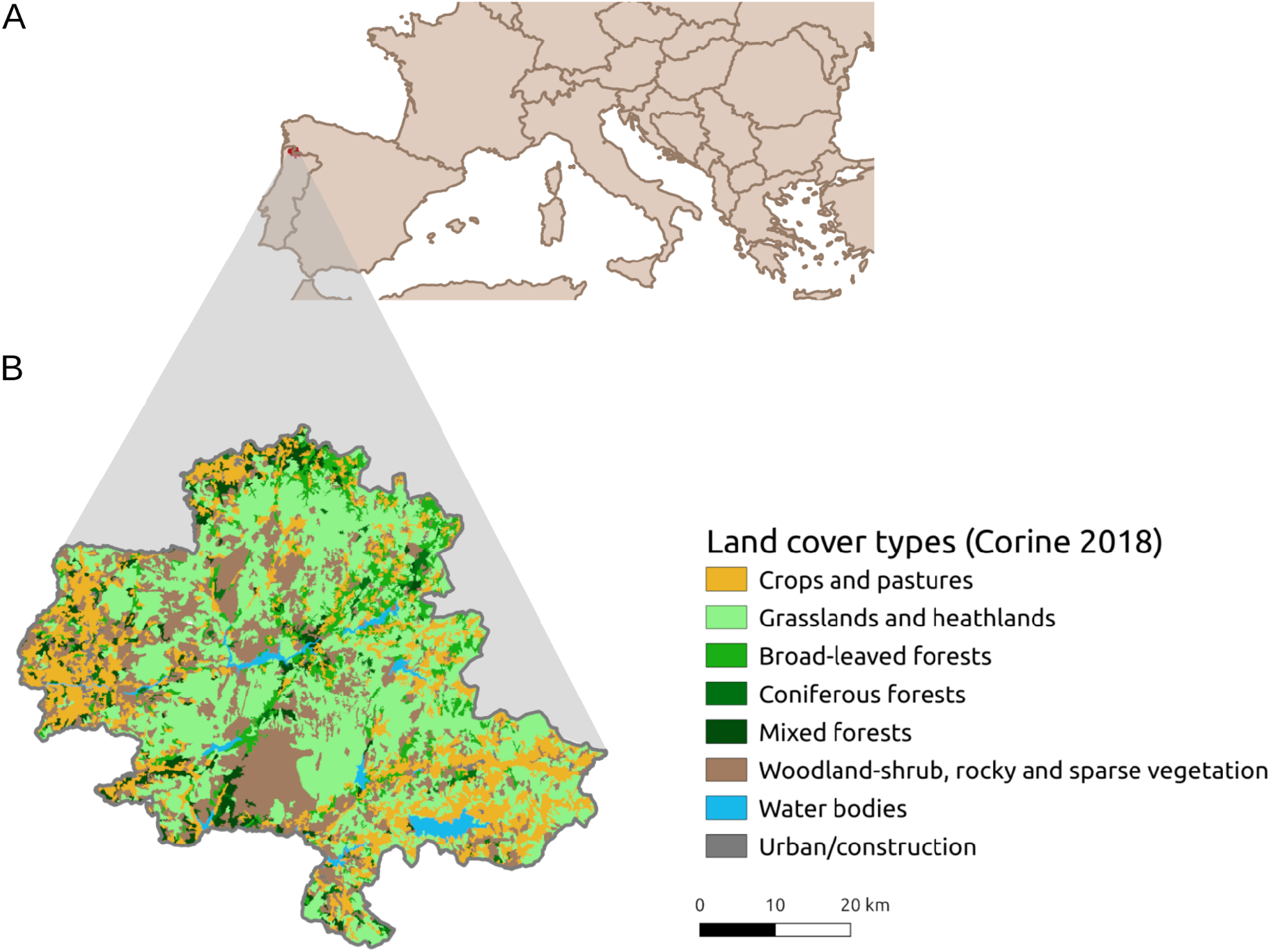
A) Location of the Gerês-Xurés Biosphere Reserve in Southern Europe and B) Land cover types of the Gerês-Xurés Biosphere Reserve based on the Corine Land cover (2018).

### 2.2. Land-use scenarios for wildfire management

Four land-use change scenarios were simulated for the study area over a 40-year period from 2010 to 2050, following Pais et al. (2020): Business as Usual (BAU), fire-smart forest conversion (fire-smart), High Nature Value farmlands (HNVf), and a combination of HNVf and fire-smart. The BAU simulates fire ignition, spread, and extinction representing the historical fire regime and land-use change trends reported between 1987 and 2010 (Pais et al., 2020; Regos et al., 2015). These historical changes are clearly dominated by farmland abandonment corresponding to 400 ha of annual conversion from cropland to shrubland, and 1.6 conversion rate from shrubland to oak due to the natural succession (Pais et al., 2020). The fire-smart scenario corresponds to a strategy that fosters more fire-resistant (less flammable) and/or fire-resilient landscapes by promoting forest-type conversion from evergreen (mostly pine plantations) to native oak forests. To do so, this scenario uses the highest conversion rate from shrubland to oak among all fire-smart scenarios evaluated by Pais et al. (2020) (i.e. 2.4, considering that 1.6 is the natural succession rate) with the historical land abandonment of 400 ha of annual conversion from cropland to shrubland and a rate of 1 of conversion from coniferous to deciduous woodlands. The HNVf scenario was developed to envisage a new CAP policy, wherein the abandonment of extensive agriculture is reversed (Moreira and Pe’er, 2018). This is, agricultural crops gradually expand over previously abandoned areas (shrublands). We selected the HNVf scenario with the highest amount of land-cover change (i.e. 1,600 annual ha from shrubland to cropland) among all scenarios in Pais et al. (2020) because it represents the best-case scenario where changes in the fire regime are more evident (i.e. burned and suppressed areas). Finally, the last scenario, HNVf + fire-smart, incorporates conversions towards more resistant and less flammable forest types (fire-smart) into new land-use policies promoting agricultural areas (HNVf). In this scenario, the highest conversion rates from shrubland to oaks among all scenarios (i.e. 2.4 conversion rate) and from shrubland to cropland (i.e. 1,600 annual ha) were applied. All scenarios assume the same level of suppression, considering the current firefighting levels based on fire exclusion policies (see Pais et al., 2020 for details).

### 2.3. Fire landscape modelling

We used a spatially explicit process-based model, REMAINS, which integrates the main factors driving fire-landscape dynamics in Southern European mountain landscapes (Pais et al., 2020). The REMAINS model considers how the spatio-temporal interactions between fire and vegetation dynamics, fire suppression, and land-use changes affect the fire regime (and consequently landscape composition and dynamics) at short and medium timescales. It reproduces fire-landscape dynamics according to the pre-designed scenario storylines explained above, simulating wildfires (including fire ignition, spread, burning, and extinction), vegetation dynamics (i.e. natural succession and post-fire regeneration), land-use changes (e.g. agriculture abandonment or intensification), and forest-type conversions (e.g. increase in intensive plantations for timber production). At each time step (1 year), the model simulated fire ignition, spread, and extinction. Fires were simulated each year until the potential annual area to be burned (i.e. the target area) was reached in each scenario. The target annual area refers to the area expected to burn according to historical fire data from 1983 to 2010 (INCF, 2020; MITECO, 2020). Since the case study area expanded over Spain and Portugal, fire size distributions were taken one per country as model inputs, and the actual final fire sizes (burnt areas) emerged from the spatial interaction between the location of fire ignitions, landscape composition and configuration, topography, and fire suppression. The probability of fire ignition is a function of human and biophysical variables (Pais et al.,2020). The spread rate was formulated as a polynomial expression with three factors (slope, aspect, and fire-proneness of each land-cover type) adapted from Duane et al. (2016). Two fire-suppression strategies are implemented: (1) ‘active fire suppression’, in which suppression of a fire front starts when the fire spread rate is below a specific threshold, mimicking the current capacity of fire brigades to extinguish low-intensity fires; and (2) ‘passive fire suppression’, based on firefighting opportunities created by the presence of agricultural areas (set as 1 ha) which break the continuity of highly flammable vegetation. Therefore, this fire-suppression strategy mimics the advantage that fire brigades can take from heterogeneous low-fuel landscape mosaics. The model tracks not only the area effectively burned in each fire event but also the target area to be burnt in the absence of suppression over the entire simulated period for all scenarios, allowing us to calculate the avoided burned area for all scenarios. The wildfire management scenarios set the changes in land-use changes and the REMAINS model uses a spatial procedure following Aquilué et al. (2017) to allocate the quantity of change (i.e. to select the cells to be transformed to the target land-cover type). The quantity of change at each time step was based on a landscape change analysis performed using historical information for the 1987-2010 period (details can be found in Pais et al. (2020a)), which was simulated in locations with a higher likelihood of being transformed to the target land-cover type, using the neighbour factor approach introduced by Verburg et al. (2004). We ran 100 replicates for each scenario to deal with the uncertainty arising from fire stochasticity (i.e. the spatial distribution of fire ignitions and subsequent spread) for the 2010-2050 period (see Pais et al. 2020 for more details).

### 2.4. Economic evaluation of net wildfire damages on ecosystem services

To compare landscape policy scenarios, we assumed that the wildfire manager would choose the alternative that generates the lowest suppression spending and wildfire damages on ecosystem services provision. Thus, we estimate the net present value of suppression costs and wildfire ecosystem services damages associated with the net changes in burned land covers in the case study area over the years simulated (we use a 40-year time horizon and a 3% discount rate, δ). Discounting future benefits is standard in economic analysis, even though it is subject to great debate (Price, 2014). The discount rate used here fits within the time preference discount rate range (0 - 8%) often applied to forestry projects (Sauter and Mußhoff, 2018). Our framework is as follows:

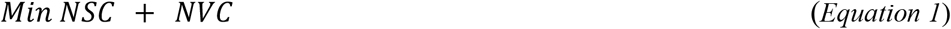

with

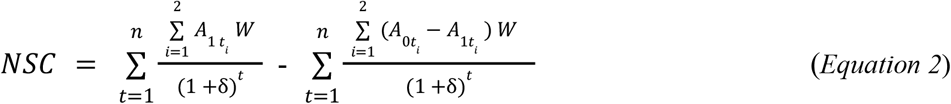

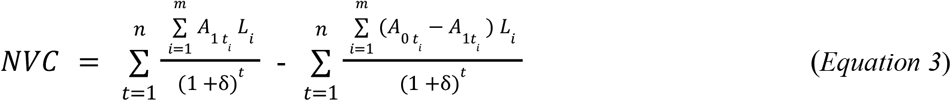

where *NSC* and *NVC* represent the present value of net suppression costs and net value change in wildfire damages, respectively. The elements contained in the NSC and NVC include the suppression spending and wildfire ecosystem services damages associated with the wildfire regime under each land-use policy (first term in equations 2 and 3, respectively), which are net of the potential beneficial effects, in terms of reduced wildfire burned area, arising from the suppression capabilities of the simulated land-use changes (the second term in equations 2 and 3, represent avoided suppression costs and wildfire damages, respectively). *A*_0*ti*_ and *A*_1*ti*_ represent the REMAINS burned area output under the baseline situation (i.e. historical trend without suppression effort), and under the land-use policy simulated for each year *t* and land cover *i*, respectively. We calculated the discounted value of net suppression costs for each fire season by multiplying the average suppression cost per hectare, W, for the annual burned and avoided burned land cover *i* of the evergreen and deciduous forests. When the *NSC* is positive it represents a cost; and when NSC is negative it represents saving from the land-use policy, and the more negative it is the larger the saving. This occurs because the avoided suppression costs due to the suppression capabilities of the landscape exceeded the suppression costs incurred over the period simulated. The suppression cost per hectare was based on Vázquez-Vázquez et al. (2014), who used wildfire reports of 6,383 fires in the Galician XV forest district, where part of our study area is located, for the period 1999-2008. Similarly, the present value of the net damage on ecosystem services for all the fire seasons simulated, depends on the number of hectares burned, the type of land cover *m* burned (croplands, shrublands, evergreen forests and deciduous forest), and avoided to be burned, and the monetary value per ha associated with the ecosystem service affected, *L*. The computation of these net wildfire damages was carried out as a collection of economic models for each of the most relevant economic activities associated with ecosystem services provision, in the Biosphere Reserve: agricultural, pasture, forestry, and recreational use. We also included wildfire impacts on climate regulation because of the role that woodland landscapes can play in supporting the European commitment to net zero. We based our analysis on market-based valuation approaches, using market prices, replacement costs, and avoided costs, as follows to quantify the damages per ha for each of the ecosystem services included in our analysis.

#### 2.4.1. Food provision

Agricultural land accounts for 8% of the total study area. Production activities are essentially dedicated to mixed crop-livestock farming, and our analysis focused on the five most abundant crops in the Gerês-Xurés: cereals for grain, fodder crops, potatoes, fruit trees, and vineyards. The annual economic returns of agricultural production were obtained as the sum of the per-hectare annual returns for crop *j*(*p_jt_Y_jt_* - *c_jt_*) multiplied by the number of hectares burned (or avoided burned) dedicated to crop *j* (H_*jt*_) as follows:

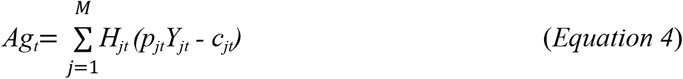

where *H_jt_* is obtained by multiplying the burned or avoided burned area of agricultural land cover obtained by the model in year *t* (*H_t_*) by the proportion of the total area dedicated to crop *j* (*w_jt_*); *p_jt_* is the observed market price; *Y_jt_* is the productivity per hectare; and *c_it_* is the cost of production per hectare. The proportion of crop *j* (*w_j_*) in the entire Gerês-Xurés area was obtained as an average of the proportion of crop*i* in Portugal (Instituto Nacional de Estatística, www.ine.pt) and Spain (Instituto Galego de Estatística, www.ige.gal) weighted by the importance of each region in the total area of the Gerês-Xurés (Table S1). Prices, productivity, and cost for each crop were obtained from official Spanish statistics, i.e. Statistical Yearbook and Farm Cost Studies (Ministerio de Agricultura, 2000-2018, 2009-2019, 2010-2017), weighted, in the case of cereals for grain, fodder crops, and fruit trees, by the specific crop percentage obtained from the 2019 Spanish Statistical Yearbook (Ministerio de Agricultura, 2009-2019; Table S2). To estimate the predicted future prices and costs of each crop (2019-2050), we applied the compound annual growth rate observed in the available period (2009-2019), using the average of half first data values as the initial value and the average of half last data values as the last value to avoid the strong variation observed in the time series. We assumed that productivity would not change over the simulation period in the study area because no significant technological improvements were expected.

#### 2.4.2. Pasture provision

The landscape in the study area is dominated by shrublands (32%). This area is used for extensive agropastoral activities and is therefore associated with pasture production (Celaya et al., 2022). However, since the agro-livestock uses in mountainous areas almost disappeared in the region in recent decades (Corbelle-Rico et al. 2022), we assumed that only 10% of the shrubland area is oriented toward animal feeding. The value of the annual economic returns of the damages (and avoided damages) on pasture production in year *t* was obtained as the product of the per-hectare annual pasture returns multiplied by the number of hectares of shrubland burned (and avoided to be burned). Prices (*p_t_*) and productivity (*Y_t_*) of the fodder crops were assigned to pasture production, using a replacement cost to farmers in fodder, and zero cost was considered as the result of natural regeneration. The compound annual growth rate observed in the available period for fodder crop prices (2009-2019) was used to estimate future pasture prices.

#### 2.4.3. Timber provision

Coniferous plantations cover 11% of the total study area and are largely used to generate benefits for timber production (Pasalodos-Tato et al., 2010). The net present value of the wildfire damages (and avoided damages) on forestry returns depends on the productivity of the parcel for growing timber (i.e. site quality), the net price of timber, rotation time, and the percentage of value destroyed due to wildfire. The land expectation value *sensu* Faustam (Amacher et al., 2009) was calculated to assess the impact of wood production on annual economic returns under simulated landscape policies. The land expectation value is a common discounted cash flow method applied to value forest stands (Tahvonen, 2004) dedicated to perpetuity to forestry (i.e. a perpetual series of rotations), and annual returns are computed as the interest on this natural capital stock using an interest rate of 3%. Based on the growth model for even-aged stands of maritime pine (*Pinus pinaster*) in Galicia proposed by González et al. (1999, 2005), we calculated the timber growth and volume. The even-aged maritime pine stand was assumed to be of site class (height of dominant trees) of 130 dm, which corresponds to the interior areas of Galicia, with an initial density of 1,300 trees/ha and a natural mortality rate of 0.01 trees/ha. Therefore, we captured the fact that inland stands have lower growth in height than coastal stands (Rodríguez-Soalleiro et al., 1994). Mean net prices/m^3^ at the roadside for pulpwood with bark were obtained from the Database prices of the Galician Forestry Association for the 1990-2021 period (Asociación Forestal de Galicia, 1990-2021), ranging from 18.2 €/m^3^ to 23 €/m^3^. Following Pasalodos-Tato et al. (2010) study in the region of the case study area, the percentage of *Pinus pinaster* timber value reduced after a wildfire was taken to be 25%. The timber price change rate over the years simulated was calculated as described in section 2.4.1. The cost of planting was assumed to be zero because forests are in general not managed in the area and we also assumed no silvicultural activities occur due to a lack of data availability.

#### 2.4.4. Recreational benefits

Recreational use is one of the most recognised ecosystem services provided by the Gerês-Xurés Biosphere Reserve, especially in Portugal, where its only national park, Peneda-Gerês National Park, is located and has a large number of visitors (Martins et al., 2021). We estimated recreation based on the presence of deciduous woodlands, mostly represented by oak forests (18% of the studied area) and old conifer forests, which are considered to be highly valued for recreation (Ciesielski and Stereńczak, 2018; Löf et al., 2016; Norman et al., 2010). A recent study also suggested that oak is the tree species most appreciated by the local population (Calviño-Cancela and Cañizo-Novelle, 2018). To estimate the monetary value of recreational benefits, we used the travel cost study of Mendes and Proença, (2009), conducted in Peneda-Gerês National Park, which generates an average recreational benefit per visitor of 194€, with values ranging from 116€ to 448€. The per-hectare recreational benefits were obtained by multiplying this average recreational value by the average annual number of visits over the years 2017-2019 obtained from the Institute for Nature Conservation and Forest (2020), accounting for 111,000 visitors per year. This resulted in a value of 297 € per hectare, which was used for the Portuguese side of the study area, and was halved in the Spanish area (149 €) due to its relatively smaller attractiveness. These values are consistent with the wide range of monetary benefits of forest recreation in Mediterranean countries (Merlo and Croitoru, 2005). The value of the annual economic returns of recreation in year *t* was obtained as the product of the per-hectare annual returns multiplied by the number of hectares burned and avoided to be burned dedicated to deciduous forest and more than 60 years old coniferous forest.

#### 2.4.5. Climate regulation

We applied the Integrated Valuation of Ecosystem Services and Tradeoffs (InVEST) Carbon Storage and Sequestration module (Sharp et al., 2020) to assess the climate regulation ecosystem service by computing the carbon sequestrated in the entire landscape. The InVEST carbon module is a spatially explicit tool that uses carbon stocks in four different pools (above- and below-ground biomass, litter, and soil organic carbon) in each land cover class to estimate carbon storage per map pixel. The amount of carbon sequestration across a particular time period (in our case a decade) was computed by comparing the carbon storage levels in each map pixel of the landscape at the beginning of the time period with that stored in the area at the end of the time period. Therefore, carbon sequestration (or emission) only occurs when a map pixel of a given land cover type changes between the beginning and end of a period, which in our case is driven by fire-vegetation and/or land cover change; otherwise, the carbon sequestration/emission rate will be zero. This analysis used the land cover map output from REMAINS for the years 2010, 2020, 2030, and 2050, while the estimates of carbon stocks for each land cover class in the study area were based on data collection (i.e. published scientific literature, official statistics from the Portuguese and Spanish national forest inventories) and/or modelling (see Pais et al. 2020 for a detailed description). We used the social cost of carbon as a standard metric to compute the costs (benefits) of emitting (sequestering) CO_2_ in policy evaluation. This represents the discounted sum of additional damages by an incremental tonne of CO_2_ emissions in a particular year. The uncertainties in Integrated Assessment Models have generated considerable variation in the available estimates of the social cost of carbon; and they have also been heavily criticised due to the highly sensitive assumptions of these models related to discount rates, damage functions, or future projected emissions (Pindyck, 2013; Stern, 2016). Wang et al. (2019) meta-analysis showed that values can range from −50 to 8.752 $/t C (13.36e2386.91 $/t CO2), with a mean value of 200.57 $/tC (54.70 $/t CO2). We took a conservative approach within this range and applied a social cost of carbon of 44 $ t C (Tol, 2008); which is also smaller than the US government’s most cited mean value of $51 per t CO2, using a 3% discount rate (Interagency Working Group on Social Cost of Greenhouse Gases, 2021). It is however higher than the market prices of voluntary carbon credits, which would reflect the average economic income that forest owners in the study area could earn by selling credits (Forest Trends’ Ecosystem Marketplace, 2021); because these market prices underestimate the social cost (benefits) of emissions (avoiding emissions) (Wegner and Pascual, 2011). The present value benefits for the climate regulation under the alternative land-use scenarios for the Gerês-Xurés over the studied period were converted to euros by applying a conversion rate of 0.8619 (average conversion rate between 9 March 2020 and 9 March 2021 - European Central Bank); and the average discounted annual benefit per ha was used to compute the damages (and avoided damages) to estimate the net value change on this ecosystem service.

## 3. Results

### 3.1. Predicted burned and avoided burned area trends for each land cover type and their associated ecosystem services

Overall, policies promoting HNVf by itself, and in combination with fire-smart forest conversions (i.e. the HNVf + fire-smart scenario) were predicted to increase the avoided burned area for all land-cover types, in particular those linked to agricultural land cover and, to a lesser extent, oak land cover (Fig. 2b). However, agricultural land was found to be more exposed to wildfire due to the gradual increase of the extent of this cover type promoted by agricultural policies (see HNVf and HNVf + fire-smart scenarios Fig. 2a). The oak native woodlands exposed to burning increases over the studied period under the large-scale fire-smart forest type conversion toward more fire resistant species (Fig. 2a). According to our simulations, land cover with coniferous trees is predicted to decrease over the simulated period under all scenarios due to progressive reduction caused by fire and the fact that scenarios do not include tree planting activities (Fig. 2). Shrublands in sparsely vegetated areas are predicted to remain constant at approximately 1,500 ha/year under the BAU and fire-smart scenarios due to natural succession processes for the coming decades (Fig. 2a). The gradual conversion of this land-cover type to agricultural lands and oak forests (under HNVf and HNVf + fire-smart scenarios, respectively) is captured through a decrease in its coverage in the study area, and therefore it is gradually less affected by fires (Fig. 2).

**Figure 2.**
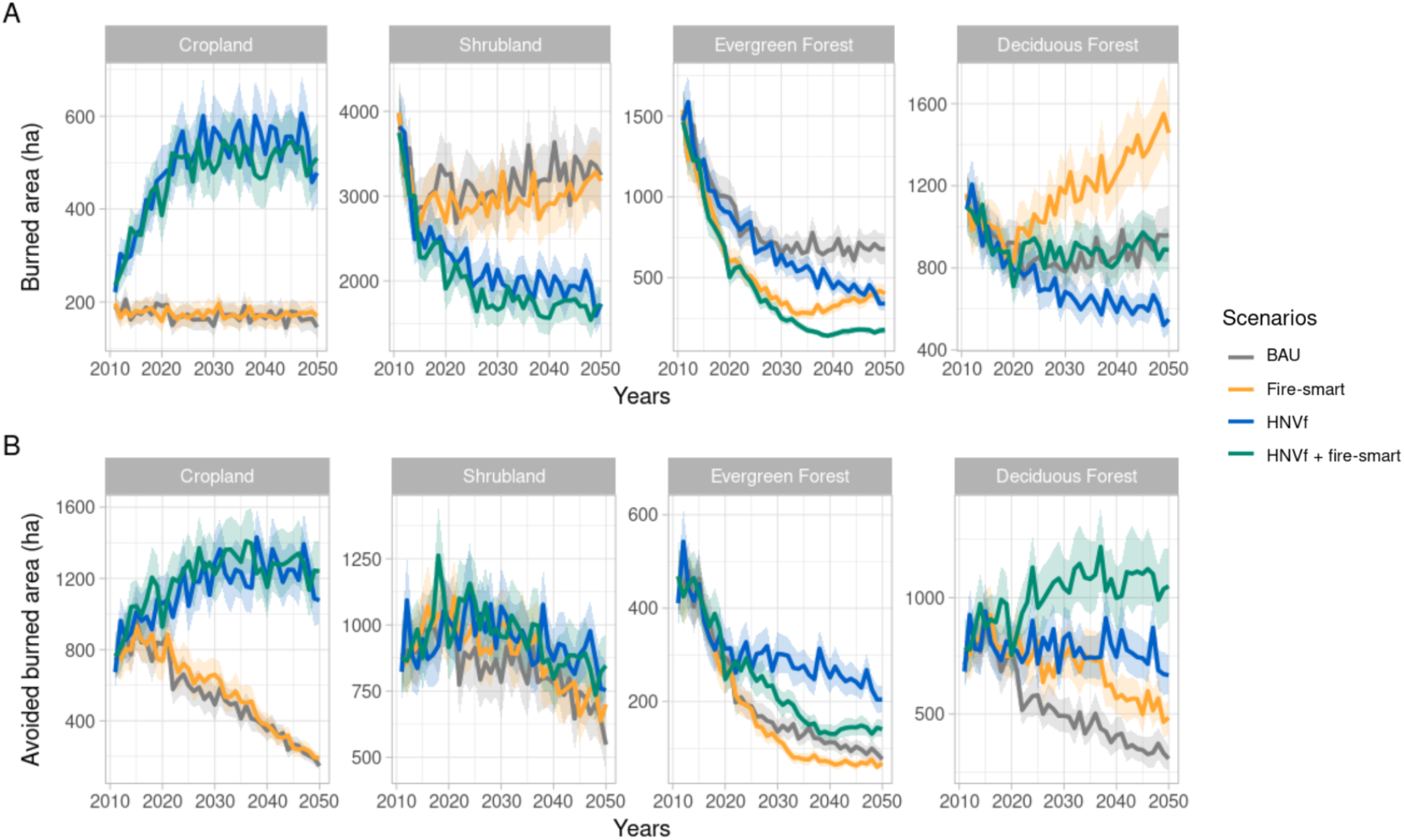
Temporal changes (2010-2050) of a) burned area (ha) and b) avoided burned area (ha) for the land-cover types associated with ecosystem services (i.e. croplands to food production, shrublands to pasture production, evergreen forest to timber production, and forests to recreation) under the land-use management scenarios (Business-as-Usual (BAU), fire-smart, High Nature Value Farmlands (HVNf), and HVNf + fire-smart). For all plots, colored lines indicate mean values while the transparent colored areas indicate the error limits defined by the mean range values.

### 3.2. Wildfire net suppression costs and ecosystem services net value change under land-use management scenarios

HNVf and its combination with more fire-resistant tree species, i.e. HNVf + fire-smart, generate the best suppression cost outcomes, which is consistent with the fact that these two scenarios generate the largest avoided burned area (32,000 ha and 34,000 ha for HNVf and HNVf+fire-smart, respectively). The results show that these scenarios generate negative net suppression costs, this is, they lead to savings in suppression costs because of the large benefits generated in terms of avoided costs (Fig. 3). The expected present value of net suppression costs is −3,959 K€ for HNVf + fire-smart and −3,922 K€ for HNVf (Table S3). In contrast, the expected present value of net suppression cost in the fire-smart scenario and BAU is positive, and therefore represents a cost, which it is higher in the fire-smart scenario (2,308 K€) than that in the BAU scenario (1,149 K€) (Table S3), as a result of having a larger burned area of deciduous trees than in the BAU (Fig. 2).

**Figure 3.**
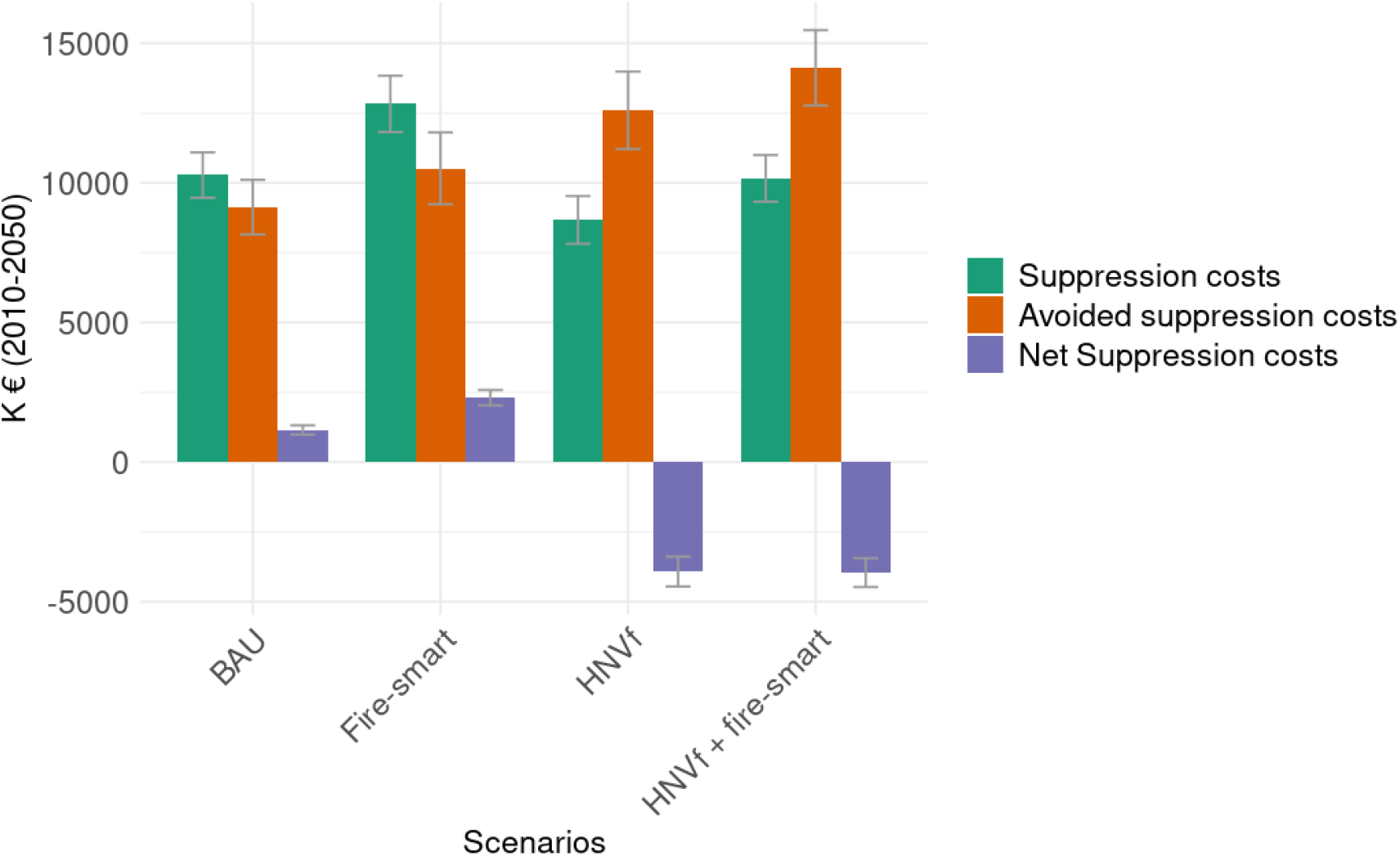
Present value of wildfire suppression costs, avoided suppression costs and net suppression costs under land-use management scenarios (Business-as-Usual (BAU), fire-smart, High Nature Value Farmlands (HVNf), and HVNf + fire-smart) over the 50 years simulated period.

The largest component of the cost-plus-net-value change is the wildfire net damages on ecosystem services flows. Fig. 4a and 4b show the mean present value of wildfire damages and avoided damages from reduced burned area, respectively. All scenarios show the small expected private returns associated with crop production; which is due to the fact that costs are expected to rise at a higher rate than prices over the period analysed. Therefore, under all scenarios, the net present value of damages in crop production are very modest (ranging from 958 K€ for HNVf to 1,269 K€ for BAU; see Table S4), when valued purely on market prices, as farmers’ financial returns that are lost due to wildfire. These results reflect the existing trade-offs under HNVf policies between the financial returns to crop producers in the study area and the societal gains from fire suppression savings. The wildfire net damages are slightly lower for pasture production in all scenarios because of the assumption that only a very small fraction of shrubland is under this type of use. In fact, the largest expected damages in private returns of fires are associated with timber production, mainly under the BAU and HNVf scenarios, as these generate the most burned are of coniferous forests. Promoting more fire-resistant species (fire-smart and HNVf + fire-smart) leads to a significant reduction in the damages and avoided damages on private forest returns from timber production compared to the BAU scenario (52% and 25% smaller in terms of net damages, respectively). The net present value of societal benefits from climate regulation over the period studied under HNVf land use policies is very limited (12,408 K€); therefore, it is not surprising that the damages (fig. 4a) and avoided damages (fig. 4b) on this ecosystem service are practically negligible. The highest net present value from carbon sequestration and storage is under the fire-smart strategy (375,998 K€). Fire-smart also generates the highest recreational values, and thus the highest wildfire damages on this ecosystem service (fig. 4a, Table S4) because of the higher presence of deciduous forests in relation to other scenarios. Combining both HNVf and fire-smart policies generates the largest avoided damages on these recreational benefits (6,255 K€, Table S4). Finally, our results show that the lowest societal discounted net suppression costs and ecosystem services damages are associated with the HNVf + fire-smart scenario with 7,132 K€ for the period 2010-2050 (fig. 5, Table S5). This is because this scenario results in suppression cost savings from agricultural expansion, while also generating a significant reduction in damages on timber and recreational benefits. In contrast, the least efficient scenario is BAU, representing land abandonment, which generates discounted societal net costs and damages of 25,710 K€ (fig. 5, Table S5).

**Figure 4.**
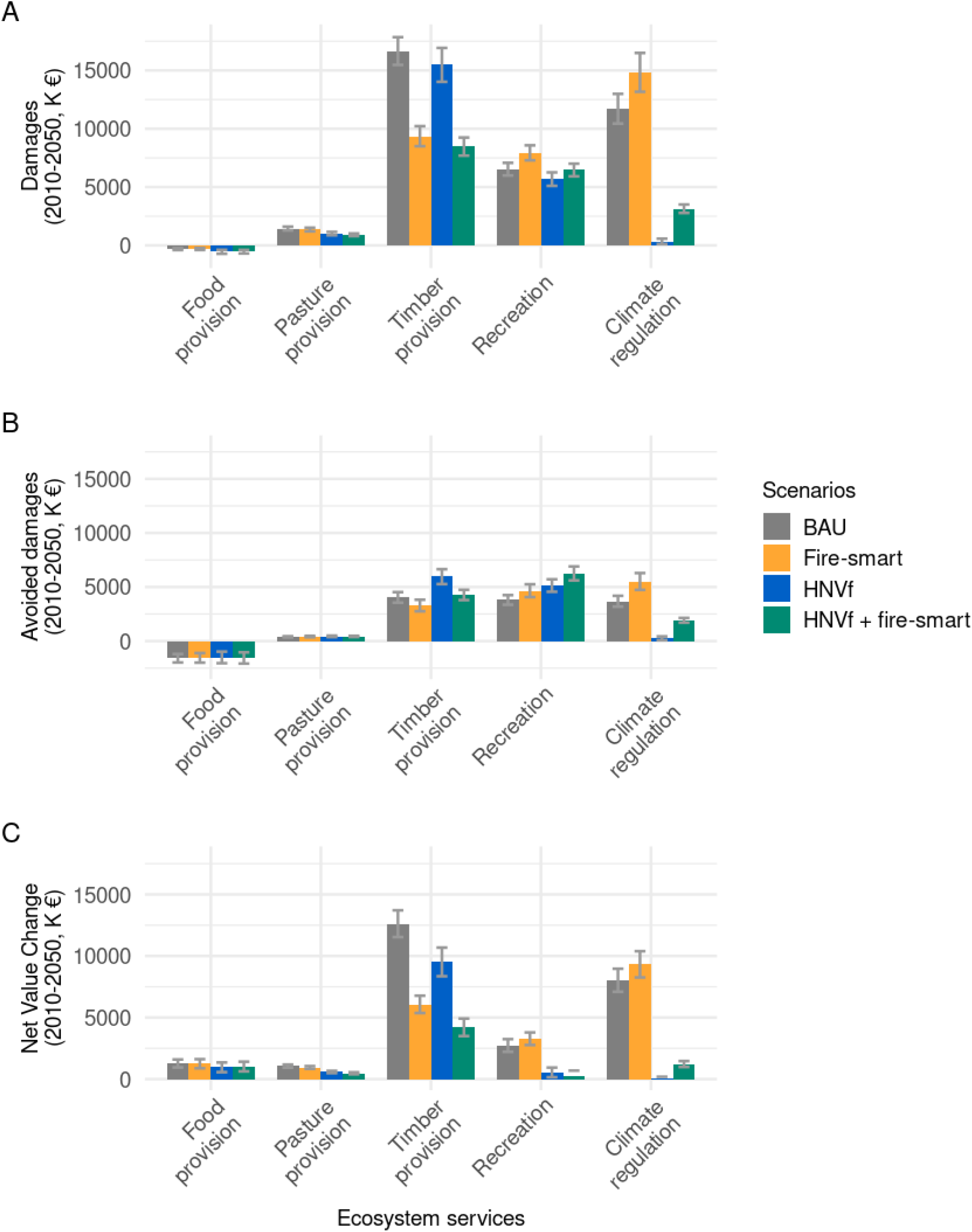
Bar plots of a) damages; b) avoided damages; and c) net value change (damages - avoided damages) for each ecosystem service (food provision, pasture provision, timber provision, recreation and climate regulation) over the 2010-2050 period under the different scenarios (Business-as-Usual (BAU), fire-smart, High Nature Value Farmlands (HVNf), and HVNf + fire-smart). Error bars indicate the standard deviation computed across the 100-fold REMAINS simulations.

**Figure 5.**
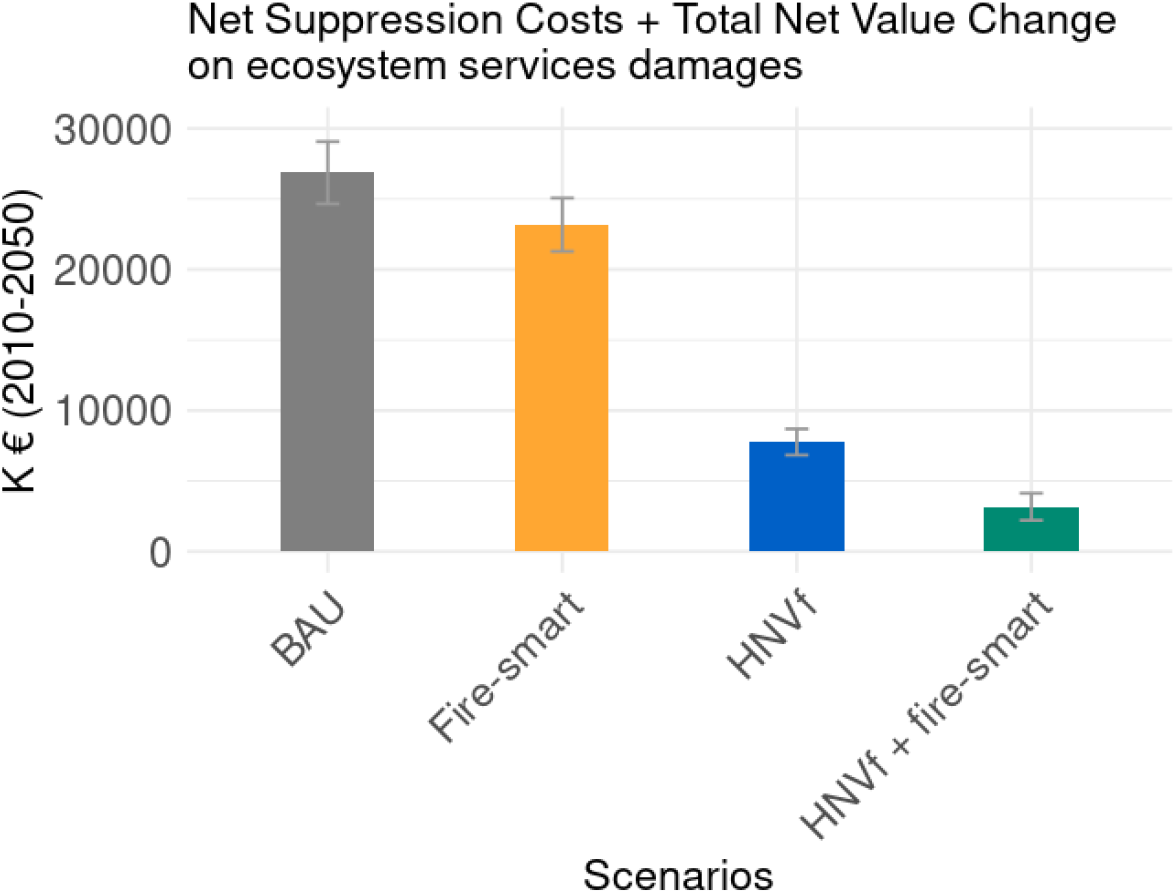
Present value of net suppression costs + total net value change on ecosystem services damages over the 2010-2050 period under the different scenarios (Business-as-Usual (BAU), fire-smart, High Nature Value Farmlands (HVNf), and HVNf + fire-smart). Error bars indicate the standard deviation computed across the 100-fold REMAINS simulations.

## 4. Discussion

Reverting land abandonment through recultivation and promoting fire-resistant tree species in woodland creation has been shown to be an attractive way to reduce wildfire hazard, to the detriment of more flammable landscapes such as shrublands (Aquilué et al., 2020; Moreira and Pe’er, 2018; Pais et al., 2020). We applied the least-cost-plus-net-value-change approach of wildland fire economics and estimated net changes in wildfire damages based on their implications for ecosystem services that affect financial returns to landowners in the study area (e.g. agriculture, pasture, and timber), and wider economic benefits (recreation and climate regulation). Changes in the flow of these ecosystem services are simulated under alternative land-use scenarios that affect wildfire regimes and vegetation dynamics outcomes (including fire ignition, spread, extinction or post-fire regeneration) over a 40-year time horizon. Thus, we computed the expected present value of suppression and wildfire costs and avoided costs of the land use management options to inform wildfire decision-making. The Transboundary Biosphere Reserve Gerês-Xurés (Spain-Portugal) was used as an illustrative case of the abandoned rural landscapes of Southern Europe.

Firstly, our results suggest that the potential net suppression cost savings can be substantial under the HNVf-related scenarios. The expected present value of suppression cost savings is 3,959 K€ for HNVf + fire-smart and 3,922 K€ for HNVf land use policies. Therefore, our study highlights the effects of HNVf agricultural policies in reducing governmental suppression of wildfire costs. This is consistent with previous studies that also concluded that the promotion of extensive agriculture increases avoided burned area in agricultural and forest land covers; and promoting agriculture can help suppress wildfires to the detriment of more flammable landscapes such as shrublands (Aquilué et al., 2020; Moreira and Pe’er, 2018; Pais et al., 2020). This is also consistent with other studies in the same area that suggested that large-scale forest conversions to more fire-resistant forests (fire-smart scenario) would not be enough to effectively reduce potential burned area (Pais et al., 2020), a strategy that would be effective in a matrix of agricultural areas. Our results are in line with existing literature suggesting that farmland abandonment would decrease the fire regulation capacity and the fire protection ecosystem services in mountain landscapes (see e.g. Sil et al 2019). In this regard, our simulations also suggest that if agricultural abandonment continues over the next few decades (see BAU and fire-smart scenarios in Fig. 4), the avoided fire damages on the targeted ecosystem services would likely be lower compared to scenarios that promote agriculture (see HNVf and HNVf + fire-smart in Fig. 4).

Secondly, we found that the most efficient scenario that generates the lowest expected present value of suppression costs plus NVC, is HNVf + fire-smart, whereas the worst scenario is BAU. However, the lack of financial viability of agriculture shown under all scenarios in our results made the implementation of HNVf policies to address wildfire management challenging. In fact, agriculture abandonment has been predominant in the study area as well as in Southern Europe in the last decades, consequently increasing wildfire risk (Estoque et al., 2019; Mantero et al., 2020; Terres et al., 2015). Thus, our findings suggest that as long as policies fail to cope with agricultural abandonment, this wildfire risk is expected to persist; as farmers are only financially rewarded for commodity production, but not for the provision of ecosystem services and their contribution to suppressing wildfires. These findings call for the need for ecosystem services accounting into agricultural policies and develop policies that encourage farmers’ choices that enhance the social benefits from crop and pasture production, including their role on fire suppression shown here. In this sense, payments for ecosystem services (Rodríguez-Ortega et al., 2018) could enhance the maintenance of farmland landscapes that would contribute to fire regulation (Sil et al., 2019). Even though designing policies to improve the efficiency of these policy instruments is not a simple task (Börner et al., 2017; Engel, 2016), our results are consistent with studies in the case study area that found significant beneficial effects of public policies that incentivise recultivation of abandoned farmlands for agriculture and animal husbandry (Corbelle-Rico et al., 2022).

Thirdly, the trade-offs among various policy strategies highlight the importance of undertaking an ecosystem service approach when assessing wildfire land-use policies, since all potential land-use policies affect ecosystem functioning and its ability to deliver a wide range of ecosystem services in fire-prone systems (Raviv et al., 2021). Even though HNVf was the best strategy for suppression cost savings, we found that it generated the lowest expected present value for climate regulation because carbon sequestration in farmlands is very limited. In fact, the combination of HNVf with fire-smart seems critical because fire-smart complements HNVf by introducing fire-resistant and resilient tree species that sequester carbon and thus regulate climate. In addition, HNVf + fire-smart added co-benefits in reducing timber damages and increasing recreational benefits (Fig. 4). Therefore, future land-use policies should not only enhance HNVf to reduce wildfire impacts (and suppression costs reduction) but should also complement them by promoting fire-resistant and resilient tree species to avoid losing the landscape societal benefits associated with climate regulation. Moreover, previous studies in the area have predicted that BAU would overall decline species habitat suitability and that HNVf and fire-smart policies could be beneficial for biodiversity conservation (Pais et al., 2020). In particular, promoting agriculture would benefit bird species breeding in open habitats, whereas HNVf + fire-smart could benefit reptile species by providing more favourable habitats for thermoregulation, shelter and food availability (Pais et al., 2020).

We have assessed the damages and avoided damages on ecosystem services resulting from alternative landscape planning as a wildfire management policy, yet some challenges remain. Our results are likely to underestimate the economic impact of wildfires in the future because climate change was not explicitly included in the scenarios within the fire landscape model. Future studies should therefore include climate-fire relationships and interactions among climate, vegetation dynamics, and fire management (Abatzoglou et al., 2018). Information on fire severity could also complement the study by providing a more accurate estimate of what is lost (e.g. high fire severity may involve more negative impacts than low severity). Future work could also include a wide range of ecosystem services impacts. For example, we only included wildfire impacts on recreational benefits due to a lack of data availability and the qualitative nature of other cultural values (e.g. sense of place, identity and spiritual value). Including the economic valuation of regulating ecosystem services beyond climate regulation, such as those related to habitat quality or biodiversity, could also provide a more accurate picture given the relevance of nature conservation within the Biosphere Reserve case study area (e.g. Campos et al., 2021). A further extension of this study could also explicitly address the spatial arrangements of the proposed land-use changes, which can facilitate answering relevant management questions, such as how the promoted land-use changes can be arrayed across the landscape to achieve the greatest reduction in wildfire economic losses. This could address our simple assumption regarding fire suppression costs with a single value per hectare for all study areas, which can be considered unrealistic since spatial factors could affect suppression costs (e.g. proximity and accessibility to inhabited areas). Similarly, the agricultural pasture and forestry yield were assumed to be the same across all landscapes (as in other studies such as Butry et al. (2010) and Mercer et al. (2007)), ignoring that some areas could be more productive than others. Finally, further research is needed to estimate the full least cost-plus-net-value-change model that includes estimates of the implementation costs of the different land-use policies investigated here, to provide better tools for assessing prevention policies on wildfire management.

## 5. Conclusions

Our results added economic evidence to recent research about the critical role that fire-smart agroforestry policies could play to promote sustainable solutions to the wildfire problem in abandoned rural landscapes of Southern Europe (Aquilué et al., 2020; Moreira and Pe’er, 2018; Pais et al., 2020). Promoting extensive agriculture is shown here to provide fire-suppression opportunities, generating societal benefits in the form of savings in fire suppression costs. However, the effect on suppression costs must be weighted against the effect on ecosystem services from these landscape changes as wildfire strategies. To address this, we constructed an estimate of the net-value-change in ecosystem services flows under the simulated landscape policies. Our results showed that large-scale forest conversion to more fire-resistant trees would not be on their own the most economically effective solution to reduce potential burned area and consequently suppression costs; however, when integrated with HNVf policies to jointly reduce fire hazards, this strategy generates the lowest net suppression cost and wildfire ecosystem services damages. In this sense, the new European Common Agricultural Policy offers an excellent opportunity to incorporate fire-smartness into renewed EU agricultural policies that would contribute to wildfire costs and damage mitigation. Our findings emphasise the need for design payments for ecosystem services as a governance approach to reward private landowners’ services for the wildfire protection of their crops. However, while a large amount of strategically allocated cropland areas (at least 1,200 ha per year in the Biosphere Reserve ‘Gêres-Xurés’ in our simulations) could be gradually incorporated into the landscape over the next few decades to significantly reduce the suppression cost from wildfires; we also showed that relying on promoting crop could reduce public good climate regulation services from the landscape. This emphasises the need to fully evaluate the ecosystem services trade-offs in wildfire management decision-making.

## Acknowledgements

This research was supported by Portuguese national funds through FCT - Foundation for Science and Technology, I.P., under the FirESmart project (PCIF/MOG/0083/2017). J.L-D is funded by the Alexander von Humboldt Foundation. A.R. is funded by the Spanish Ministry of Science and Innovation (IJC2019-041033-I). N.A. is funded by the Spanish Ministry of Science and Innovation (FCJ2020-046387-I). Â. Sil received support from the Portuguese Foundation for Science and Technology (FCT) through Ph.D. Grant SFRH/BD/132838/2017, funded by the Ministry of Science, Technology and Higher Education, and by the European Social Fund - Operational Program Human Capital within the 2014-2020 EU Strategic Framework. This project has contributed to the European Horizon 2020 research and innovation programme under grant agreement No 101037419 (FIRE-RES project).

## Author contributions

Conceptualization: AR, JT, MC-A; Data curation: JL-D, MC-A, JT, NA, ÂS; Formal analysis: JL-D; Funding acquisition: AR; Methodology: JT, MC-A; Supervision: JT, MC-A, AR; Writing: JT, JL-D, MC-A, AR. All authors review and comment on the manuscript before submission.

## Appendix A. Supplementary data

**Table S1.** Percentage of the area of each type of crop in the Gerês-Xurés Portuguese and Spanish sectors. Total area refers to the percentage of each crop weighted by the contribution of each country to the total extent of the Biosphere Reserve (i.e. 71% in Portugal and 29% in Spain).

|  | Area in the Gerês-Xurés (in %) |  | Total area weighted (in %) |
| --- | --- | --- | --- |
|  | Portugal | Spain |  |
| Cereals for grain | 39.0 | 65.4 | 46.7 |
| Fodder crops | 39.8 | 9.9 | 31.1 |
| Potatoes | 6.1 | 18.6 | 9.7 |
| Fruit trees | 2.8 | 2.4 | 2.7 |
| Vineyards | 12.3 | 3.7 | 9.8 |
| Total | 100.0 | 100.0 | 100.0 |

**Table S2.**
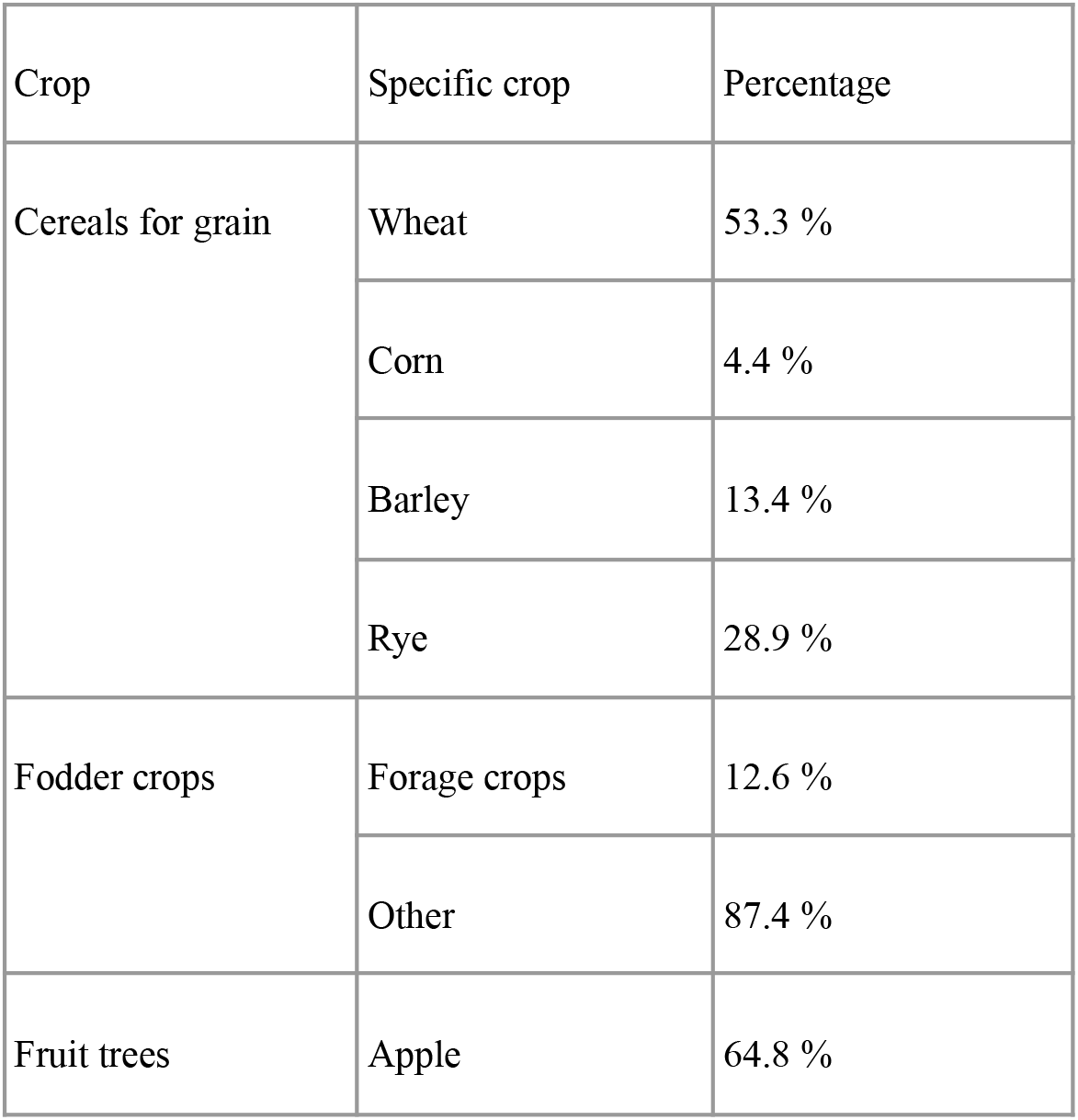
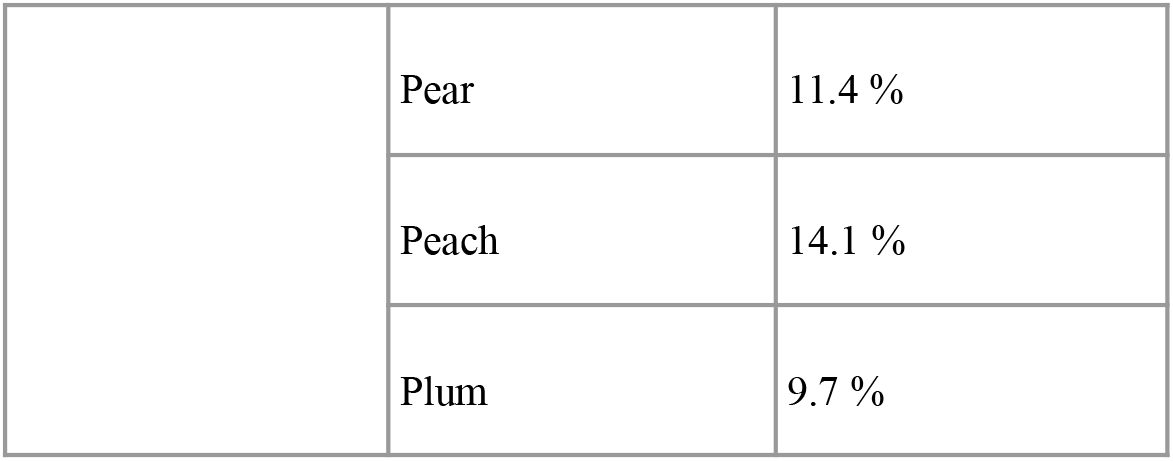
Percentage of surface for specific crops of cereals for grain, fodder crops, and fruit trees.

**Table S3.** Average and standard deviation of suppression costs, avoided suppression costs and net suppression costs over the 2010-2050 period under the different scenarios (Business-as-Usual (BAU), fire-smart, High Nature Value Farmlands (HVNf), and HVNf + fire-smart).

| Scenario | Suppression costs |  | Avoided suppression costs |  | Net suppression costs |  |
| --- | --- | --- | --- | --- | --- | --- |
|  | Mean (K€) | Standard deviation (K€) | Mean (K€) | Standard deviation (K€) | Mean (K€) | Standard deviation (K€) |
| BAU | 10282 | 813 | 9133 | 979 | 1149 | -166 |
| Fire-smart | 12830 | 1010 | 10522 | 1286 | 2308 | -275 |
| HNVf | 8677 | 854 | 12599 | 1388 | -3922 | -534 |
| HNVf + fire-smart | 10164 | 836 | 14123 | 1350 | -3959 | -514 |

**Table S4.** Average and standard deviation of damages, avoided damages and net value change (damages - avoided damages) for each ecosystem service (food provision, pasture provision, timber provision, recreation and climate regulation) over the 2010-2050 period under the different scenarios (Business-as-Usual (BAU), fire-smart, High Nature Value Farmlands (HVNf), and HVNf + fire-smart).

| Scenario | Ecosystem Services | Damages |  | Avoided damages |  | Net Value Change |  |
| --- | --- | --- | --- | --- | --- | --- | --- |
|  |  | Mean (K€) | Standard deviation (K€) | Mean (K€) | Standard deviation (K€) | Mean (K€) | Standard deviation (K€) |
| BAU | Food provision | -321 | 80 | -1590 | 391 | 1269 | 323 |
|  | Pasture provision | 1447 | 160 | 389 | 58 | 1058 | 125 |
|  | Timber provision | 16660 | 1188 | 4039 | 488 | 12621 | 1087 |
|  | Recreation | 6533 | 541 | 3803 | 443 | 2730 | 514 |
|  | Climate regulation | 11714 | 1273 | 3683 | 504 | 8031 | 935 |
| Fire-smart | Food provision | -307 | 84 | -1555 | 441 | 1248 | 370 |
|  | Pasture provision | 1358 | 158 | 413 | 63 | 945 | 110 |
|  | Timber provision | 9359 | 860 | 3288 | 534 | 6071 | 704 |
|  | Recreation | 7936 | 642 | 4657 | 593 | 3279 | 511 |
|  | Climate regulation | 14833 | 1664 | 5511 | 778 | 9322 | 1065 |
| HNVf | Food provision | -553 | 170 | -1512 | 535 | 958 | 392 |
|  | Pasture provision | 1010 | 145 | 434 | 67 | 577 | 90 |
|  | Timber provision | 15476 | 1453 | 5958 | 687 | 9519 | 1164 |
|  | Recreation | 5679 | 579 | 5135 | 576 | 544 | 391 |
|  | Climate regulation | 344 | 231 | 256 | 172 | 88 | 103 |
| HNVf + fire-smart | Food provision | -543 | 153 | -1565 | 520 | 1022 | 393 |
|  | Pasture provision | 906 | 111 | 431 | 53 | 475 | 79 |
|  | Timber provision | 8469 | 777 | 4265 | 477 | 4204 | 710 |
|  | Recreation | 6462 | 543 | 6255 | 648 | 207 | 481 |
|  | Climate regulation | 3143 | 361 | 1919 | 218 | 1224 | 240 |

**Table S5.** Average and standard deviation of total Net Value Change and Net Suppression Costs + total Net Value Change over the 2010-2050 period under the different scenarios (Business-as-Usual (BAU), fire-smart, High Nature Value Farmlands (HVNf), and HVNf + fire-smart).

| Scenario | Total Net Value Change |  | Net Suppression Costs + Total Net Value Change |  |
| --- | --- | --- | --- | --- |
|  | Mean (K€) | Standard deviation (K€) | Mean (K€) | Standard deviation (K€) |
| BAU | 25710 | 2372 | 26858 | 2206 |
| Fire-smart | 20865 | 2166 | 23173 | 1890 |
| HNVf | 11686 | 1462 | 7763 | 929 |
| HNVf + fire-smart | 7132 | 1472 | 3173 | 958 |

## References

Abatzoglou, J.T., Williams, A.P., 2016. Impact of anthropogenic climate change on wildfire across western US forests. Proc. Natl. Acad. Sci. 113, 11770–11775.

Abatzoglou, J.T., Williams, A.P., Boschetti, L., Zubkova, M., Kolden, C.A., 2018. Global patterns of interannual climate–fire relationships. Glob. Change Biol. 24, 5164–5175. https://doi.org/10.1111/gcb.14405

Alcasena, F.J., Salis, M., Nauslar, N.J., Aguinaga, A.E., Vega-García, C., 2016. Quantifying economic losses from wildfires in black pine afforestations of northern Spain. For. Policy Econ. 73, 153–167. https://doi.org/10.1016/j.forpol.2016.09.005

Amacher, G.S., Ollikainen, M., Koskela, E., 2009. Economics of Forest Resources. The MIT Press, Cambridge, Massachusetts.

Aquilué, N., De Cáceres, M., Fortin, M.J., Fall, A., Brotons, L., 2017. A spatial allocation procedure to model land-use/land-cover changes: Accounting for occurrence and spread processes. Ecol. Model. 344, 73–86. https://doi.org/10.1016/j.ecolmodel.2016.11.005

Aquilué, N., Fortin, M.J., Messier, C., Brotons, L., 2020. The Potential of Agricultural Conversion to Shape Forest Fire Regimes in Mediterranean Landscapes. Ecosystems 23, 34–51. https://doi.org/10.1007/s10021-019-00385-7

Asociación Forestal de Galicia, 1990. Prezos orientativos da madeira galega.

Börner, J., Baylis, K., Corbera, E., Ezzine-de-Blas, D., Honey-Rosés, J., Persson, U.M., Wunder, S., 2017. The Effectiveness of Payments for Environmental Services. World Dev. 96, 359–374.

Butry, D.T., Prestemon, J.P., 2019. Economics of WUI/Wildfire Prevention and Education, in:Manzello, S.L. (Ed.), Encyclopedia of Wildfires and Wildland-Urban Interface (WUI) Fires. Springer International Publishing, Cham, pp. 1–8. https://doi.org/10.1007/978-3-319-51727-8_105-1

Butry, D.T., Prestemon, J.P., Abt, K.L., Sutphen, R., Butry, D.T., Prestemon, J.P., Abt, K.L., Sutphen, R., 2010. Economic optimisation of wildfire intervention activities. Int. J. Wildland Fire 19, 659–672. https://doi.org/10.1071/WF09090

Calviño-Cancela, M., Cañizo-Novelle, N., 2018. Human dimensions of wildfires in NW Spain:Causes, value of the burned vegetation and administrative measures. PeerJ 2018. https://doi.org/10.7717/peerj.5657

Calviño-Cancela, M., Chas-Amil, M.L., García-Martínez, E.D., Touza, J., 2016. Wildfire risk associated with different vegetation types within and outside wildland-urban interfaces. For.Ecol. Manag. 372, 1–9. https://doi.org/10.1016/j.foreco.2016.04.002

Campos, F.S., David, J., Lourenço-de-Moraes, R., Rodrigues, P., Silva, B., Vieira da Silva, C., Cabral, P., 2021. The economic and ecological benefits of saving ecosystems to protect services. J.Clean. Prod. 311, 127551. https://doi.org/10.1016/j.jclepro.2021.127551

Castellnou, M., Prat-Guitart, N., Arilla, E., Larrañaga, A., Nebot, E., Castellarnau, X., Vendrell, J., Pallàs, J., Herrera, J., Monturiol, M., Cespedes, J., Pagès, J., Gallardo, C., Miralles, M., 2019. Empowering strategic decision-making for wildfire management: avoiding the fear trap and creating a resilient landscape. Fire Ecol. 15, 31, s42408-019-0048–6. https://doi.org/10.1186/s42408-019-0048-6

Celaya, R., Ferreira, L.M.M., Lorenzo, J.M., Echegaray, N., Crecente, S., Serrano, E., Busqué, J., 2022. Livestock Management for the Delivery of Ecosystem Services in Fire-Prone Shrublands of Atlantic Iberia. Sustainability 14, 2775. https://doi.org/10.3390/su14052775

Chas-Amil, M.L., Prestemon, J.P., McClean, C.J., Touza, J., 2015. Human-ignited wildfire patterns and responses to policy shifts. Appl. Geogr. 56, 164–176. https://doi.org/10.1016/j.apgeog.2014.11.025

Chas-Amil, M.L., Touza, J., Prestemon, J.P., 2010. Spatial distribution of human-caused forest fires in Galicia (NW Spain). WIT Trans. Ecol. Environ. 137, 247–258. https://doi.org/10.2495/FIVA100221

Ciesielski, M., Stereńczak, K., 2018. What do we expect from forests? The European view of public demands. J. Environ. Manage. 209, 139–151. https://doi.org/10.1016/j.jenvman.2017.12.032

Corbelle-Rico, E., Sánchez-Fernández, P., López-Iglesias, E., Lago-Peñas, S., Da-Rocha, J.-M., 2022. Putting land to work: An evaluation of the economic effects of recultivating abandoned farmland. Land Use Policy 112, 105808. https://doi.org/10.1016/j.landusepol.2021.105808

Donovan, G.H., Rideout, D.B., 2003. A Reformulation of the Cost Plus Net Value Change (C+NVC)Model of Wildfire Economics. For. Sci. 6.

Duane, A., Aquilué, N., Gil-Tena, A., Brotons, L., 2016. Integrating fire spread patterns in fire modelling at landscape scale. Environ. Model. Softw. 86, 219–231. https://doi.org/10.1016/j.envsoft.2016.10.001

Elia, M., Lovreglio, R., Ranieri, N.A., Sanesi, G., Lafortezza, R., 2016. Cost-effectiveness of fuel removals in mediterraneanwildland-urban interfaces threatened by wildfires. Forests 7, 1–11. https://doi.org/10.3390/f7070149

Engel, S., 2016. The Devil in the Detail: A Practical Guide on Designing Payments for Environmental Services. Int. Rev. Environ. Resour. Econ. 9, 131–177. https://doi.org/10.1561/101.00000076

Estoque, R.C., Gomi, K., Togawa, T., Ooba, M., Hijioka, Y., Akiyama, C.M., Nakamura, S., Yoshioka, A., Kuroda, K., 2019. Scenario-based land abandonment projections: Method,application and implications. Sci. Total Environ. 692, 903–916. https://doi.org/10.1016/j.scitotenv.2019.07.204

Fernandes, P.M., 2022. Make Europe’s forests climate-smart and fire-smart. Nature 609, 32–32. https://doi.org/10.1038/d41586-022-02318-2

Fernandes, P.M., 2013. Fire-smart management of forest landscapes in the Mediterranean basin under global change. Landsc. Urban Plan. 110, 175–182. https://doi.org/10.1016/j.landurbplan.2012.10.014

Florec, V., Burton, M., Pannell, D., Kelso, J., Milne, G., 2020. Where to prescribe burn: the costs and benefits of prescribed burning close to houses. Int. J. Wildland Fire 29, 440. https://doi.org/10.1071/WF18192

Forest Trends’ Ecosystem Marketplace, 2021. State of Forest Carbon Finance 2021. Washington DC:Forest Trends Association.

González, J.Á.-C., Soalleiro, R.R., Alonso, G.V., 1999. Elaboración de un modelo de crecimiento dinámico para rodales regulares de “Pinus pinaster Ait” en Galicia. For. Syst. 8, 319–334.

González, J.G.Á., González, A.D.R., Soalleiro, R.R., Anta, M.B., 2005. Ecoregional site index models for Pinus pinaster in Galicia (northwestern Spain). Ann. For. Sci. 62, 115–127. https://doi.org/10.1051/forest:2005003

Hesseln, H., 2000. The Economics of Prescribed Burning: A Research Review. For. Sci. 46, 322–334.

Houtman, R.M., Montgomery, C.A., Gagnon, A.R., Calkin, D.E., Dietterich, T.G., McGregor, S., Crowley, M., Houtman, R.M., Montgomery, C.A., Gagnon, A.R., Calkin, D.E., Dietterich, T.G., McGregor, S., Crowley, M., 2013. Allowing a wildfire to burn: estimating the effect on future fire suppression costs. Int. J. Wildland Fire 22, 871–882. https://doi.org/10.1071/WF12157

INCF, 2020. Incêndios Rurais [WWW Document]. URL http://www2.icnf.pt/portal/florestas/dfci/inc (accessed 10.25.21).

Institute for Nature Conservation and Forest, 2020. Visitor’s at the Nature Areas-Portugal.

Interagency Working Group on Social Cost of Greenhouse Gases, U.S.G., 2021. Technical Support Document: Social Cost of Carbon, Methane, and Nitrous Oxide Interim Estimates under Executive Order 13990 48.

IPCC, 2018. Global Warming of 1.5°C. An IPCC Special Report on the impacts of global warming of 1.5°C above pre-industrial levels and related global greenhouse gas emission pathways, in the context of strengthening the global response to the threat of climate change,.

Kottek, M., Grieser, J., Beck, C., Rudolf, B., Rubel, F., 2006. World Map of the Koppen-Geiger climate classification updated. Meteorol. Z. 15, 259–263.

Lecina-Diaz, J., Campos, J., Pais, S., Carvalho-Santos, C., Azevedo, J., Fernandes, P., Gonçalves, J., Aquilué, N., Roces-Díaz, J., Torre, M.A. de la, Brotons, L., Chas-Amil, M.-L., Lomba, A.,Duane, A., Moreira, F., Touza, J., Hermoso, V., Sil, Â., Vicente, J., Honrado, J., Regos, A. (under review). Stakeholder perceptions of wildfire management strategies as Nature-based solutions in two Iberian Biosphere Reserves.

Löf, M., Brunet, J., Filyushkina, A., Lindbladh, M., Skovsgaard, J.P., Felton, A., 2016. Management of oak forests: striking a balance between timber production, biodiversity and cultural services. Int. J. Biodivers. Sci. Ecosyst. Serv. Manag. 12, 59–73. https://doi.org/10.1080/21513732.2015.1120780

Lomba, A., Alves, P., Jongman, R.H.G., Mccracken, D.I., 2015. Reconciling nature conservation and traditional farming practices: A spatially explicit framework to assess the extent of High Nature Value farmlands in the European countryside. Ecol. Evol. 5, 1031–1044. https://doi.org/10.1002/ece3.1415

Lydersen, J.M., Collins, B.M., Brooks, M.L., Matchett, J.R., Shive, K.L., Povak, N.A., Kane, V.R., Smith, D.F., 2017. Evidence of fuels management and fire weather influencing fire severity in an extreme fire event. Ecol. Appl. 27, 2013–2030. https://doi.org/10.1002/eap.1586

Macedo, A., Tavares, A., Fontes, A., Pinto, C., Rodrigues, Machado, C., Silva, D., Carvalho, H., Regalo, H., Osório, M., Santarém, M., Gonzalez, R., Formoso, J.C., Gonzalez, F.J., Fernández, M.A., Gil, a, Veloso, N., 2009. Propuesta para la creación de la Reserva de la Biosfera Transfronteriza Gerês/Xurés. Xunta de Galicia, Comissão de Coordenação eDesenvolvimento Regional do Norte, Instituto de Conservação da Natureza e da Biodiversidade.; Xunta de Galicia, Comissão deCoordenação e Desenvolvimento Regional do Norte, Instituto de Conservação da Natureza e da Biodiversidade.

Mantero, G., Morresi, D., Marzano, R., Motta, R., Mladenoff, D.J., Garbarino, M., 2020. The influence of land abandonment on forest disturbance regimes: a global review. Landsc. Ecol. 35, 2723–2744. https://doi.org/10.1007/s10980-020-01147-w

Martins, H., 2022. Tourism in protected áreas: the exemple of Peneda-Gerês National Park (Portugal). PASOS Rev. Tur. Patrim. Cult. 20, 1113–1128. https://doi.org/10.25145/j.pasos.2022.20.075

Martins, H., Carvalho, P., Almeida, N., 2021. Destination Brand Experience: A Study Case in Touristic Context of the Peneda-Gerês National Park. Sustainability 13, 11569. https://doi.org/10.3390/su132111569

Mendes, I., Proença, I., 2009. Measuring the Social Recreation Per-Day Net Benefit of Wildlife Amenities of a National Park: A Count-Data Travel Cost Approach, School of Economics and Management, Technical University of Lisbon, Department of Economics.

Mercer, D.E., Haight, R.G., Prestemon, J.P., 2008. Analyzing Trade-Offs Between Fuels Management,Suppression, and Damages from Wildfire, in: Holmes, T.P., Prestemon, J.P., Abt, K.L. (Eds.), The Economics of Forest Disturbances: Wildfires, Storms, and Invasive Species, Forestry Sciences. Springer Netherlands, Dordrecht, pp. 247–272. https://doi.org/10.1007/978-1-4020-4370-3_13

Mercer, D.E., Prestemon, J.P., Butry, D.T., Pye, J.M., 2007. Evaluating Alternative Prescribed Burning Policies to Reduce Net Economic Damages from Wildfire. Am. J. Agric. Econ. 89, 63–77. https://doi.org/10.1111/j.1467-8276.2007.00963.x

Merlo, M., Croitoru, L., 2005. Valuing Mediterranean Forests: Towards Total Economic Value. CABI Publishing.

Ministerio de Agricultura, P. y A., n.d. Superfícies y Producciones de Cultivos. Anuarios de Estadística [WWW Document]. 2009-2018. URL https://www.mapa.gob.es/es/estadistica/temas/publicaciones/anuario-de-estadistica/ (accessed 2.24.22a).

Ministerio de Agricultura, P. y A., n.d. Estudios de costes de explotaciones agrícolas. [WWW Document]. 2010-2017. URL https://www.mapa.gob.es/es/ministerio/servicios/analisis-y-prospectiva/ECREA_Informes-Agricolas.aspx (accessed 10.15.21b).

Ministerio de Agricultura, P. y A., n.d. Precios medios anuales de las tierras de uso agrario [WWW Document]. 2000-2018. URL https://www.mapa.gob.es/es/estadistica/temas/estadisticas-agrarias/economia/encuesta-precios-tierra/ (accessed 2.24.22c).

MITECO, 2020. Estadísticas de Incendios Forestales [WWW Document]. URL https://www.miteco.gob.es/es/biodiversidad/estadisticas/Incendios_default.aspx (accessed 10.25.21).

Moreira, F., Ascoli, D., Safford, H., Adams, M.A., Moreno, J.M., Pereira, J.M.C., Catry, F.X., Armesto, J., Bond, W., González, M.E., Curt, T., Koutsias, N., McCaw, L., Price, O., Pausas, J.G., Rigolot, E., Stephens, S., Tavsanoglu, C., Vallejo, V.R., Van Wilgen, B.W., Xanthopoulos, G., Fernandes, P.M., 2020. Wildfire management in Mediterranean-type regions: Paradigm change needed. Environ. Res. Lett. 15. https://doi.org/10.1088/1748-9326/ab541e

Moreira, F., Pe’er, G., 2018. Agricultural policy can reduce wildfires. Science 359.

Moreira, F., Viedma, O., Arianoutsou, M., Curt, T., Koutsias, N., Rigolot, E., Barbati, A., Corona, P., Vaz, P., Xanthopoulos, G., Mouillot, F., Bilgili, E., 2011. Landscape - wildfire interactions in southern Europe: Implications for landscape management. J. Environ. Manage. 92, 2389–2402. https://doi.org/10.1016/j.jenvman.2011.06.028

Norman, J., Ellingson, L., Boman, M., Mattsson, L., 2010. The value of forests for outdoor recreation in southern Sweden: are broadleaved trees important 13.

Pais, S., Aquilué, N., Campos, J., Sil, Â., Marcos, B., Martínez-Freiría, F., Domínguez, J., Brotons, L., Honrado, J.P., Regos, A., 2020. Mountain farmland protection and fire-smart management jointly reduce fire hazard and enhance biodiversity and carbon sequestration. Ecosyst. Serv. 44, 101143. https://doi.org/10.1016/j.ecoser.2020.101143

Pasalodos-Tato, M., Pukkala, T., Alboreca, A.R., Pasalodos-Tato, M., Pukkala, T., Alboreca, A.R., 2010. Optimal management of Pinus pinaster in Galicia (Spain) under risk of fire. Int. J.Wildland Fire 19, 937–948. https://doi.org/10.1071/WF08150

Pe’er, G., Bonn, A., Bruelheide, H., Dieker, P., Eisenhauer, N., Feindt, P.H., Hagedorn, G., Hansjürgens, B., Herzon, I., Lomba, Â., Marquard, E., Moreira, F., Nitsch, H., Oppermann, R., Perino, A., Röder, N., Schleyer, C., Schindler, S., Wolf, C., Zinngrebe, Y., Lakner, S., 2020. Action needed for the EU Common Agricultural Policy to address sustainability challenges. People Nat. 2, 305–316. https://doi.org/10.1002/pan3.10080

Penman, T.D., Bradstock, R.A., Price, O.F., 2014. Reducing wildfire risk to urban developments:Simulation of cost-effective fuel treatment solutions in south eastern Australia. Environ.Model. Softw. 52, 166–175. https://doi.org/10.1016/j.envsoft.2013.09.030

Pettorelli, N., Schulte to Bühne, H., Tulloch, A., Dubois, G., Macinnis-Ng, C., Queirós, A.M., Keith, D.A., Wegmann, M., Schrodt, F., Stellmes, M., Sonnenschein, R., Geller, G.N., Roy, S., Somers, B., Murray, N., Bland, L., Geijzendorffer, I., Kerr, J.T., Broszeit, S., Leitão, P.J., Duncan, C., El Serafy, G., He, K.S., Blanchard, J.L., Lucas, R., Mairota, P., Webb, T.J., Nicholson, E., 2018. Satellite remote sensing of ecosystem functions: opportunities,challenges and way forward. Remote Sens. Ecol. Conserv. 4, 71–93. https://doi.org/10.1002/rse2.59

Pindyck, R.S., 2013. Climate Change Policy: What Do the Models Tell Us? J. Econ. Lit. 51, 860–872. https://doi.org/10.1257/jel.51.3.860

Prestemon, J.P., Abt, K.L., Barbour, R.J., 2012. Quantifying the net economic benefits of mechanical wildfire hazard treatments on timberlands of the western United States. For. Policy Econ. 21, 44–53. https://doi.org/10.1016/j.forpol.2012.02.006

Price, C., 2014. Temporal aspects in forest economics, in: Handbook of Forest Resource Economics. Earthscan from Routledge, London, Great Britain, pp. 50–66.

Raviv, O., Zemah-Shamir, S., Izhaki, I., Lotan, A., 2021. The effect of wildfire and land-cover changes on the economic value of ecosystem services in Mount Carmel Biosphere Reserve,Israel. Ecosyst. Serv. 49, 101291. https://doi.org/10.1016/j.ecoser.2021.101291

Regos, A., 2022. Nature-based solutions in an era of mega-fires. Nature 607, 449–449. https://doi.org/10.1038/d41586-022-01955-x

Regos, A., Ninyerola, M., Moré, G., Pons, X., 2015. Linking land cover dynamics with driving forces in mountain landscape of the Northwestern Iberian Peninsula. Int. J. Appl. Earth Obs.Geoinformation 38, 1–14. https://doi.org/10.1016/j.jag.2014.11.010

Rodríguez Y Silva, F., González-Cabán, A., 2010. “SINAMI”: A tool for the economic evaluation of forest fire management programs in Mediterranean ecosystems. Int. J. Wildland Fire 19, 927–936. https://doi.org/10.1071/WF09015

Rodríguez-Ortega, T., Olaizola, A.M., Bernués, A., 2018. A novel management-based system of payments for ecosystem services for targeted agri-environmental policy. Ecosyst. Serv. 34, 74–84. https://doi.org/10.1016/j.ecoser.2018.09.007

Rodríguez-Soalleiro, R., Alvarez-González, J.G., Vega-Alonso, G., 1994. Pineiro do pais. Modelo diná mico de crecemento de masas regulares de Pinus pinaster Aiton en Galicia, Xunta de Galicia. Manuais Técnicos. ed. Santiago de Compostela.

San-Miguel-Ayanz, J., Camia, A., 2010. Forest fires, mapping the impacts of natural hazards and technological accidents in Europe. An overview of the last decade. In: EEA Technical Report No 13. European Environment Agency, Copenhagen.

San-Miguel-Ayanz, J., Moreno, J.M., Camia, A., 2013. Analysis of large fires in European Mediterranean landscapes: Lessons learned and perspectives. For. Ecol. Manag. 294, 11–22. https://doi.org/10.1016/j.foreco.2012.10.050

San-Miguel-Ayanz, J., Oom, D., Artes, T., Viegas, D.X., Fernandes, P., Faivre, N., Freire, S., Moore, P., Rego, F., Castellnou, M., 2020. Forest fires in Portugal in 2017, in: Science for Disaster Risk Management 2020: Acting Today, Protecting Tomorrow. Casajus Valles, A., Marin Ferrer, M., Poljanšek, K., Clark, I. (eds.), Luxembourg.

Sauter, P.A., Mußhoff, O., 2018. What is your discount rate? Experimental evidence of foresters’ risk and time preferences. Ann. For. Sci. 75, 10. https://doi.org/10.1007/s13595-017-0683-5

Sharp, R, Douglass, J., Wolny, S., Arkema, K., Bernhardt, J., Bierbower, W., Chaumont, N., Denu, D., Fisher, D., Glowinski, Griffin, R., Guannel, G., Guerry, A., Johnson, J., Hamel, P., Kennedy, C., Kim, C.K., Lacayo, M., Lonsdorf, E., Mandle, L., Rogers, L., Silver, J., Toft, J., Verutes, G., Vogl, A.L., Wood, S., Wyatt, K., 2020. InVEST 3.10.2.post28+ug.ga4e401c.d20220324 User’s Guide., The Natural Capital Project, Stanford University, University of Minnesota, The Nature Conservancy, and World Wildlife Fund. ed.

Sil, Â., Fernandes, P.M., Rodrigues, A.P., Alonso, J.M., Honrado, J.P., Perera, A., Azevedo, J.C., 2019. Farmland abandonment decreases the fire regulation capacity and the fire protection ecosystem service in mountain landscapes. Ecosyst. Serv. 36, 100908. https://doi.org/10.1016/j.ecoser.2019.100908

Stern, N., 2016. Economics: Current climate models are grossly misleading. Nature 530, 407–409. https://doi.org/10.1038/530407a

Tahvonen, O., 2004. Timber production versus old-growth preservation with endogenous prices and forest age-classes. Can. J. For. Res. 34, 1296–1310. https://doi.org/10.1139/x04-006

Terres, J.M., Scacchiafichi, L.N., Wania, A., Ambar, M., Anguiano, E., Buckwell, A., Coppola, A., Gocht, A., Källström, H.N., Pointereau, P., Strijker, D., Visek, L., Vranken, L., Zobena, A., 2015. Farmland abandonment in Europe: Identification of drivers and indicators, and development of a composite indicator of risk. Land Use Policy 49, 20–34. https://doi.org/10.1016/j.landusepol.2015.06.009

Tol, R.S.J., 2008. The Social Cost of Carbon: Trends, Outliers and Catastrophes. Economics 2, 20080025. https://doi.org/10.5018/economics-ejournal.ja.2008-25

Vázquez Vázquez, M.C., Chas Amil, M.L., Touza, J.M., 2014. Estimación de los costes de las operaciones de extinción de incendios forestales: estudio de caso en el distrito forestal de A Limia. Rev. Galega Econ. 23, 99–114.

Verburg, P.H., de Nijs, T.C.M., van Eck, J.R., Visser, H., de Jong, K., 2004. A method to analyse neighbourhood characteristics of land use patterns. Comput. Environ. Urban Syst. 28, 667–690. https://doi.org/10.1016/j.compenvurbsys.2003.07.001

Verkerk, P.J., Martinez de Arano, I., Palahí, M., 2018. The bio-economy as an opportunity to tackle wildfires in Mediterranean forest ecosystems. For. Policy Econ. 86, 1–3. https://doi.org/10.1016/j.forpol.2017.10.016

Wang, P., Deng, X., Zhou, H., Yu, S., 2019. Estimates of the social cost of carbon: A review based on meta-analysis. J. Clean. Prod. 209, 1494–1507. https://doi.org/10.1016/j.jclepro.2018.11.058

Wegner, G., Pascual, U., 2011. Cost-benefit analysis in the context of ecosystem services for human well-being: A multidisciplinary critique. Glob. Environ. Change-Hum. Policy Dimens. - Glob.Env. CHANGE 21, 492–504. https://doi.org/10.1016/j.gloenvcha.2010.12.008

